# Dynamic protein aggregation regulates bacterial dormancy depth critical for antibiotic tolerance

**DOI:** 10.1101/233890

**Authors:** Yingying Pu, Yingxing Li, Qi Ma, Tian Tian, Ziyi Zhao, Alexander F. McVey, Fan Bai

## Abstract

The ability of some bacteria within a population to tolerate antibiotic treatment is often attributed to prolonged bacterial infection^1-3^. Unlike antibiotic resistance, which generally results from genetic mutations or plasmid transfer^4,5^, antibiotic tolerance usually refers to the phenomenon that a subgroup of cells can survive high dose antibiotic treatment as a result of phenotypic heterogeneity^6,7^. Previous studies mainly associate antibiotic tolerance with cell dormancy, by hypothesizing that the lethal effects of antibiotics are disabled due to the extremely slow metabolic and proliferation rates in dormant bacteria ^8,9^. However, less is known about how surviving bacteria subsequently escape from the dormant state and resuscitate, which is equally important for disease recurrence. Here we monitored the process of bacterial antibiotic tolerance and regrowth at the single-cell level, and found that each individual survival cell shows different ‘dormancy depth’, which in return regulates whether and when it can resume growth after removal of antibiotic. The persister cells are considered to be in shallow dormancy depth, while the viable but non-culturable cells (VBNC cells) are in deep dormancy depth. We further implemented time-lapse fluorescent imaging and biochemical analysis to establish that dynamic endogenous protein aggregation is an important indicator of bacterial dormancy depth. For cells to leave the dormant state and resuscitate, clearance of cellular protein aggregates and recovery of proteostasis are required. Through additional mutagenesis studies, we found the ability to recruit functional DnaK-ClpB machineries, which facilitate protein disaggregation in an ATP-dependent manner, determines the timeline (whether and when) for bacterial regrowth. Better understanding of the key factors regulating bacterial regrowth after surviving antibiotic attack could lead to new therapeutic strategies for combating bacterial antibiotic tolerance.

---

In our experiment, overnight culture of *Escherichia coli* was exposed to a high concentration of ampicillin (150 μg ml^-1^) treatment, 20 times the minimum inhibitory concentration (MIC), and the antibiotic killing process was monitored under a microscope. Most cells were killed by lysis, however, a small group of cells, typically composed of non-proliferative dormant cells, survived 6 hours of antibiotic treatment. We continued to observe the resuscitation course of these survival cells after removing antibiotics. Some cells resumed growth very soon, meeting the classic definition of persisters (referred to as persister-fast-recovery, persister-FR, Fig. 1a top, Supplementary Video 1). In contrast, a fair portion of survival cells remained in a dormant state showing no signs of growth. When extending the observation time, some of these dormant cells escaped from dormancy and started to proliferate (referred to as persister-slow-recovery, persister-SR, Fig. 1a middle, Supplementary Video 2). The remainder of the survival cells remained dormant even after three days of observation (viable but non-culturable cells, VBNC cells^10^, Fig. 1a bottom). This result was reproducible at a population level. After we treated a bacterial population with 150 μg ml^-1^ of ampicillin for 5 hours in a tube, we cultured the survival cells on fresh LB agar plates. Alongside colonies that were visible after one day, more colonies (an increase of approximately 25%) gradually emerged over subsequent days (Fig. 1c). These results reveal that drug tolerant cells are highly heterogeneous in their ability to recover from the dormant state.

**Figure 1.**
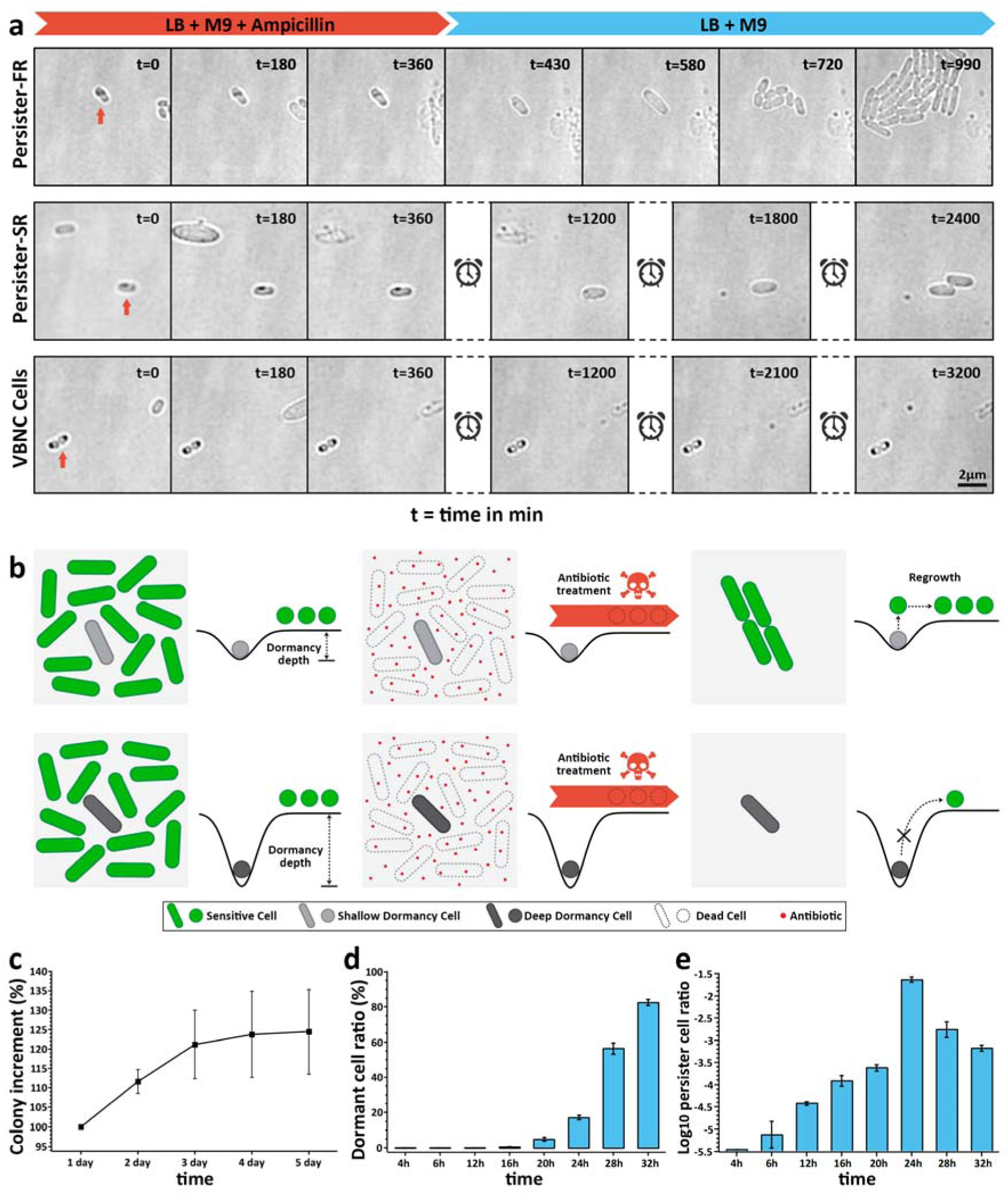
Phenomenon of bacterial dormancy is critical for antibiotic tolerance, and dormancy depth determines whether and when a cell can resuscitate. a, Timelapse images of persister-FR (upper), persister-SR (middle) and VBNC (lower) cells during antibiotic killing and subsequent resuscitation of survival cells (Supplementary Video 1). Persister-FR cells resuscitate very soon after removal of antibiotic whereas persister-SR cells resuscitate slowly. VBNC cells survive antibiotic treatment but show no sign of resuscitation in the 3 days after removal of antibiotic. Scale bar, 2 μm; t=time (min). b, Model summarising how bacterial dormancy depth regulates antibiotic tolerance and regrowth. c, Number of CFU persister cells present at indicated culture time after removal of antibiotic. d, Ratio of dormant cells as function of culture time from inoculation. Dormant cells are defined as cells non-proliferating for at least 6 hours on fresh agarose when observed under a microscope. e, Frequency of persister formation as function of culture time from inoculation. All error bars are standard errors on the mean.

Based on these observations, we postulate that the degree to which drug tolerant cells are dormant can be measured by a parameter we term ‘dormancy depth’. Cells trapped in a dormant state are protected from antibiotic killing, but after removal of antibiotic the trapped cells are able to escape from their dormant state and re-enter the active cell cycle at a rate determined by the dormancy depth. We further hypothesize that bacterial dormancy is not a single homogeneous state but displays a dormancy depth continuum. Persister-FR cells are trapped at a shallow dormancy depth and it is therefore easy for them to escape and resume growth (Fig. 1b); VBNC cells, residing at deep dormancy depths, have too large a barrier to overcome in order to escape, and thus cannot initiate resuscitation (Fig. 1b); persister-SR cells are in an intermediate state between these two extremes.

The dormancy depth appears conceptually similar to ‘quiescence depth’ in eukaryotic cells, which can be induced by serum starvation^11,12^. To probe this similarity, we tested whether bacterial dormancy depth can be modulated by culturing a bacterial sample for longer in stationary phase at 37°C. As seen in Fig. 1d, the number of dormant cells, defined as cells non-proliferating for at least 6 hours in fresh medium when observed under a microscope, increases along with inoculation time under stationary phase, indicating that longer inoculation drives more bacterial cells into deeper dormancy depth. We also performed antibiotic killing and persister counting in parallel, for bacterial populations cultured for different inoculation times. In contrast to the dormant cell ratio, the persister cell ratio initially increases steadily, peaks after 24 hours of inoculation, before decreasing gradually indicating that more dormant cells become VBNC cells (Fig. 1e, Extended Data Fig. 1). These results reveal that bacterial dormancy depth can be modulated, and in accordance with our model, the persister ratio can be optimized by increasing dormant cells in a population, whilst avoiding driving them into deep dormancy depth. Combining these results, we describe dormancy as a heterogeneous physiology state with differing depth. Bacterial drug tolerance is regulated by cell dormancy depth: as dormancy shields cells from antibiotic killing effects, dormancy depth determines whether and how survival cells resuscitate.

We then investigated what phenotypic trait can indicate bacterial ‘dormancy depth’ at a single-cell level. Under brightfield microscope observation, we noticed that a novel phenotypic feature - dark foci - often accompanied dormant cells, but was not present in actively growing cells (red arrows Fig. 1a). Normally one or two dark foci were observable per dormant cell, predominantly at the cell poles, but also in mid- or quarter-cell position. We suspected these dark foci would be protein aggregates. Thus, to test this hypothesis, we labelled IbpA, a small molecular chaperone that tightly associates with bacterial protein aggregates^13,14^, with enhanced green fluorescent protein (EGFP). As shown in Fig. 2a, IbpA-EGFP co-localises with the dark foci visible in brightfield, implying that dark foci are composed of protein aggregates.

**Figure 2.**
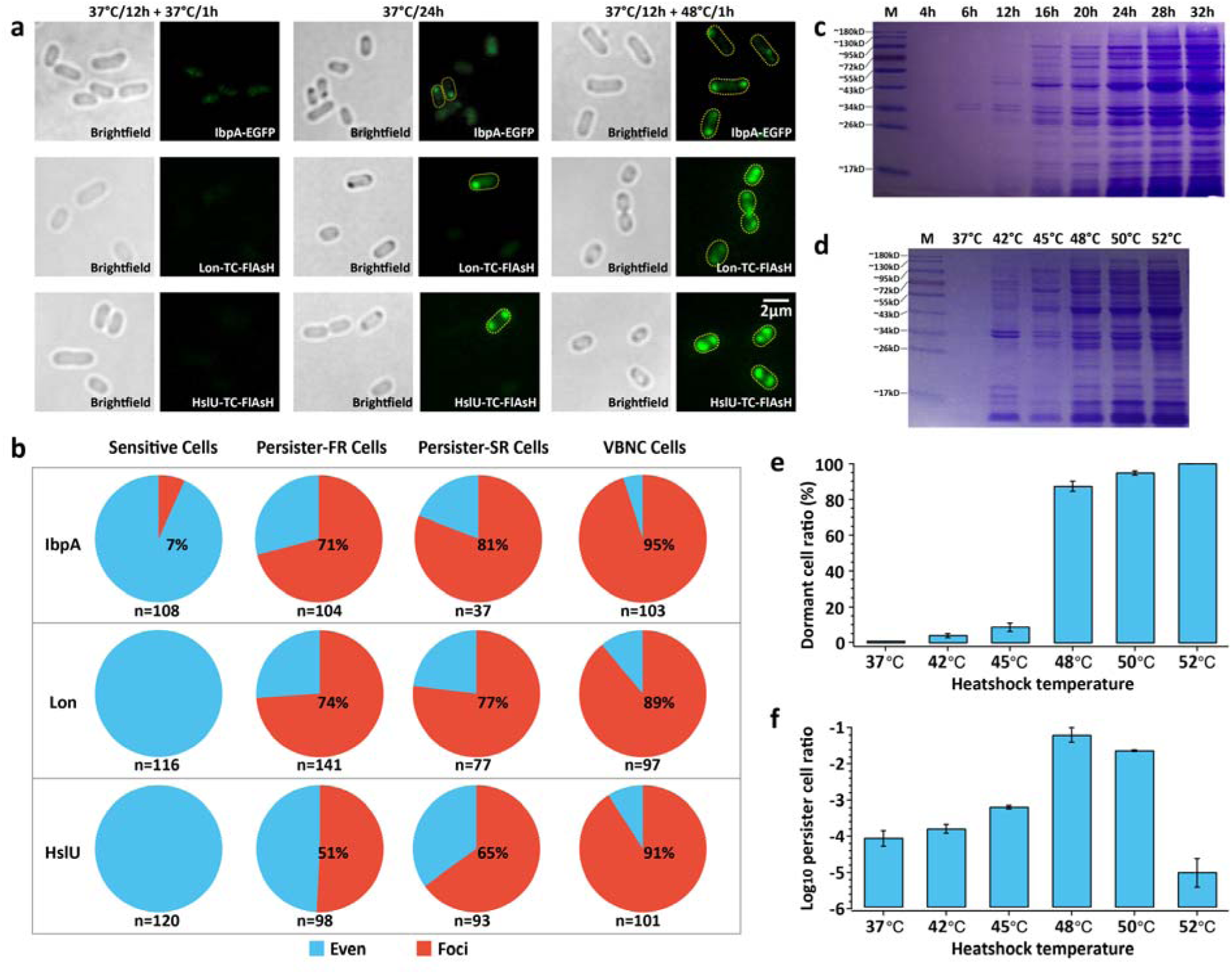
Protein aggregation induces bacterial dormancy depth. a, Brightfield and fluorescence images of cells showing protein aggregates are induced in late stationary phase or by heat shock. Protein aggregates labelled by IbpA ⸬ EGFP (upper), Lon ⸬ TC-FlAsH (middle) and HslU ⸬ TC-FlAsH (lower) (Scale bar, 2 μm). b, Percentage of cells with protein aggregates in different cell subgroups using the three fluorescent labelling methods. c-d, Fraction of endogenous insoluble protein in the cultures from indicated culture time (c) and after heat shock (d). e, Dormant cell ratio after heat shock treatment. f, Persister formation frequency after heat shock treatment, determined by antibiotic susceptibility measurement. All error bars are standard errors on the mean.

Consistent with this observation, as dormancy depth is increased by culturing cells for longer at stationary phase, the fraction of insoluble protein in the cell also increases (Fig. 2c). This further confirms that dormancy depth is strongly associated with protein aggregation. In addition, total protein mass-spectrometric analysis of 16 hour, 24 hour and 36 hour cultures identified 399, 907 and 904 types of insoluble protein respectively (Supplementary Table 1). These insoluble proteins include numerous essential and non-essential proteins involved in various important biological processes (Extended Data Fig. 3). Although the number of insoluble protein types in the 24 and 36 hour cultures is similar, analysis reveals that for almost all protein types, the insoluble fraction increases as a function of time, adding further evidence that bacteria enter deeper dormancy under prolonged stationary phase culturing.

To further test this assumption, we identified two proteins, Lon and HslU, which accumulated gradually in the insoluble protein fraction with increased culture time (Extended Table 1). In sensitive cells, these two proteins exhibit a homogeneous distribution through the cytoplasm^15,16^. We labelled these with TC-FlAsH, a marker whose fluorescence increases more than 3-fold when aggregation occurs^17^ (Extended Data Fig. 4). As expected, bright Lon-TC-FlAsH and HslU-TC-FlAsH foci co-localised with dark foci in the brightfield image, further establishing that the dark foci are formed by protein aggregates.

Next, we investigated how bacterial dormancy depth associates with protein aggregation. According to a cells response to antibiotic treatment, we divide the bacterial cell population into four subgroups: sensitive, persister-FR, persister-SR and VBNC cells characterized by a deepening dormancy depth. Within the four subgroups, the ratio of cells with protein aggregate foci, either labelled with IbpA-EGFP, Lon-TC-FlAsH or HslU-TC-FlAsH, increases with dormancy depth (Fig. 2b), indicating a strong correlation between dormancy depth and protein aggregation. The reason that IbpA is more sensitive to protein aggregates may come from its non-specificity; IbpA associates with most unfolded proteins, whereas TC-FlAsH associates with only one.

To further test if protein aggregation alone is sufficient to cause cell dormancy, we induced protein aggregation in cells by heat shock^18^ (Fig. 2d). The fraction of insoluble protein increased with increasing heat shock temperature. The corresponding dormant cell ratio also increased with cellular protein aggregation (Fig. 2e). In contrast, when heat shock temperature is below 48°C, the persister ratio of the population increased with temperature, but above this temperature the persister ratio decreased gradually (Fig. 2f), again suggesting that survival cells with deeper dormancy depth are more difficult to resuscitate. These results reinforce our dormancy depth model and suggest that cellular protein aggregation is sufficient to induce cell dormancy.

So far we have established that protein aggregation is a strong indicator of bacterial cell dormancy depth but have not examined how a cell escapes from the dormant state and resuscitates. We observed that the dark foci vanish before dormant cells re-enter the growth cycle, but persist in VBNC cells (Fig. 1a). This phenomenon suggests that protein disaggregation is a critical step for cells to escape dormancy depth and initiate resuscitation, a process analogous to restoration of proteostasis immediately prior to fertilization in *Caenorhabditis elegans* oocytes^19^. In *E. coli*, DnaK and ClpB, are responsible for dissolving protein aggregates, with DnaK binding directly to the aggregate and recruiting the disaggregase ClpB^20,21^. In the absence of DnaK, ClpB has been shown to have very low disaggregation activity; but after directly binding with DnaK, the DnaK-ClpB complex shows high efficiency of disaggregation^22^. We therefore propose that the DnaK-ClpB bichaperone system also plays a key role in dissolving protein aggregates to facilitate the regrowth of dormant cells.

To test this hypothesis, we constructed a dual-labelled strain, where Lon was labelled with TC-FlAsH and DnaK with blue fluorescent protein (BFP), to observe the dynamics of bacterial protein aggregation and disaggregation during antibiotic treatment and resuscitation. As shown in Fig. 3a-c, dormant cells that exhibit both a high intensity aggregation and dark foci survived 6 hours of high dosage antibiotic treatment, as anticipated. Surprisingly, in persister cells we found DnaK was recruited to the aggregate while antibiotic treatment was still ongoing. Upon removal of the antibiotic, TC-FlAsH fluorescent foci that were packed with DnaK molecules were diminished and cells started to regrow (Fig. 3a, Supplementary Video 3). In stark contrast, in persister-SR cells the protein level of DnaK was initially low with no obvious aggregate-colocalization pattern, and the cell showed no morphological changes. However, as with the persister-FR cells, prior to removal of the antibiotic, cellular DnaK levels increased substantially, co-localizing with protein aggregates. Upon removal of antibiotics, these protein aggregates dissolved and cell regrowth was initiated on a far longer time scale than in persister-FR cells (Fig. 3b, Supplementary Video 4). In the case of VBNC cells, the cellular DnaK level was consistently lower than either persister subgroup and no distinct aggregate-colocalisation pattern was visible. Additionally, the protein aggregates and cell morphology showed no obvious transformation during three days of observation.

**Figure 3.**
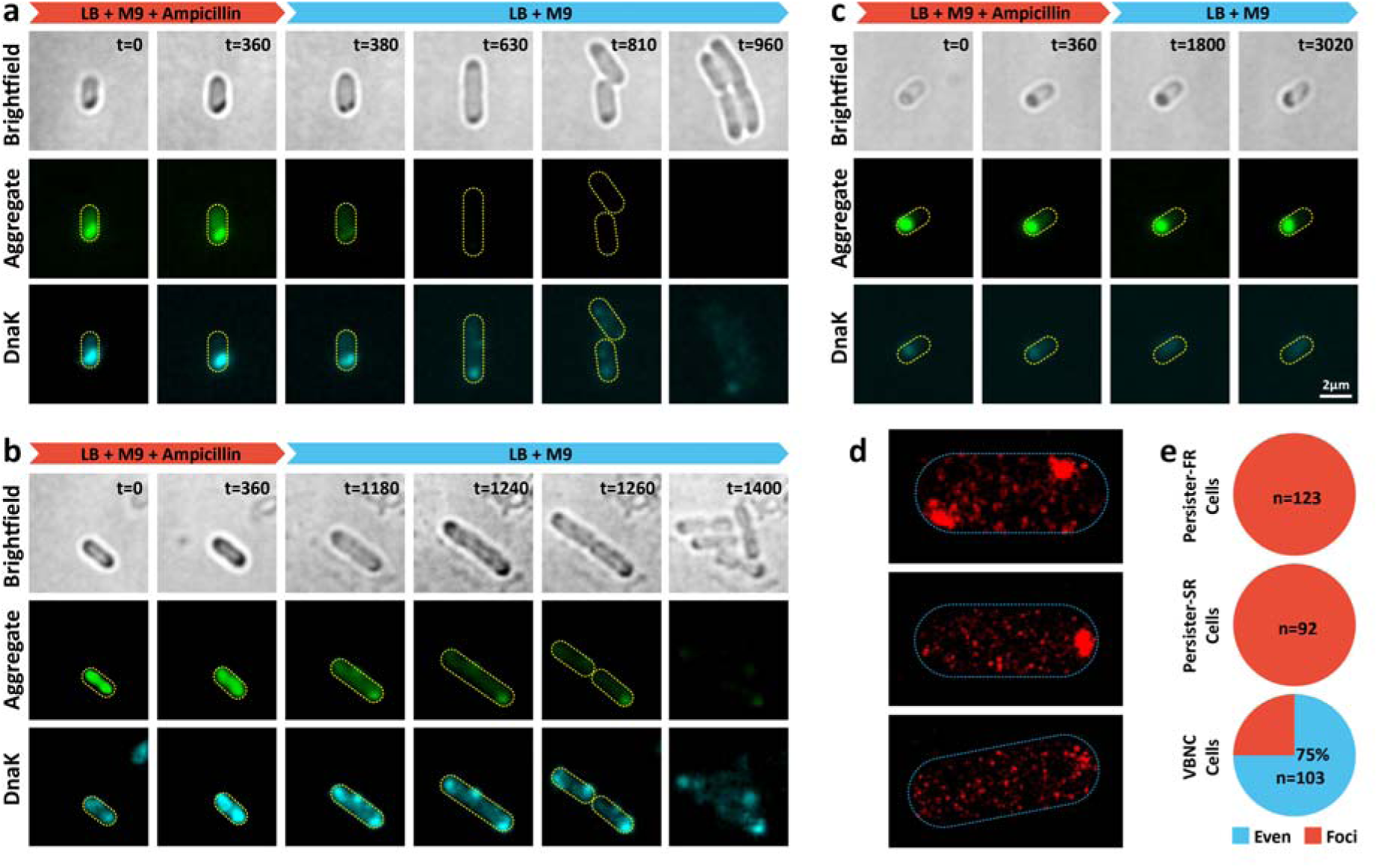
Clearance of protein aggregates and reset of cellular proteostasis before persister cell resuscitation. a-b, Time-lapse brightfield and fluorescence images showing protein aggregates and DnaK ⸬ BFP foci diminish before persister-FR (a) / persister-SR (b) cell resuscitation. c, Time-lapse brightfield and fluorescence images showing that protein aggregates remained in VBNC cells, and no visible DnaK ⸬ BFP foci was formed during our observation time. Protein aggregate is visible as dark foci in brightfield images and through Lon ⸬ TC-FlAsH foci in fluorescent images (Scale bar, 2 μm; t=time (min). d, Super-resolution images of DnaK ⸬ mMaple3. e, Percentage of cells with DnaK ⸬ BFP foci in different cell subgroups.

To confirm the foci pattern of DnaK in cells, we labelled DnaK with the photoconvertible fluorescent protein mMaple3^23^ for superresolution imaging. As shown in Fig. 3d, DnaK can either form distinct foci, mostly located at cell poles, or distribute evenly throughout the cell. In order to predict the fate of an individual cell, we traced the distribution of DnaK during exposure to the antibiotic and grouped the results according to either even or foci distributions. Our results show that DnaK foci are more prevalent (and stable) in both persister-SR and persister-FR subgroups than in VBNC or sensitive cells (Fig. 3e) with only about one-quarter of VBNC cells having a transient DnaK foci. These results show that protein disaggregation is critical for dormant cells to initiate efficient resuscitation and DnaK is an indispensable component in this process.

From our observations, we hypothesize a model to elucidate the disaggregation process and function in cell resuscitation (Fig. 4a). For a cell in protein aggregation induced dormancy, disaggregation is a prerequisite to start resuscitation. Thus, in a nutrient-rich environment such as our experiments, DnaK in the cytoplasm co-localises with protein aggregates before ClpB is also recruited through direct interaction with DnaK to initiate disaggregation. Upon disaggregation, the dormant cell is able to resuscitate and grow. To test the assumption that protein disaggregation is crucial for cell resuscitation, we constructed two knockout mutants, *ΔdnaK* and *ΔclpB*. As shown in Fig. 4b, in comparison with the wild type strain, the dormant cell rate of *ΔdnaK* and *ΔclpB* increases considerably (Fig. 4c). Normally with increased dormant cell rate one would expect an increased rate of cell survival after antibiotic treatment as well as an increased persister cell rate. However, we see an increased rate of cell survival but a decreased persister cell rate, especially in the *ΔdnaK* strain, where the resuscitation rate is extremely low (Fig. 4d). These results indicate that dormant cells with deficient disaggregation apparatus, regardless of shallow or deep dormancy depth, struggle to escape the dormancy trap. This deficient apparatus assists the cells in circumventing the disruptive effect of antibiotics, but also prevents them from resuscitation.

**Figure 4.**
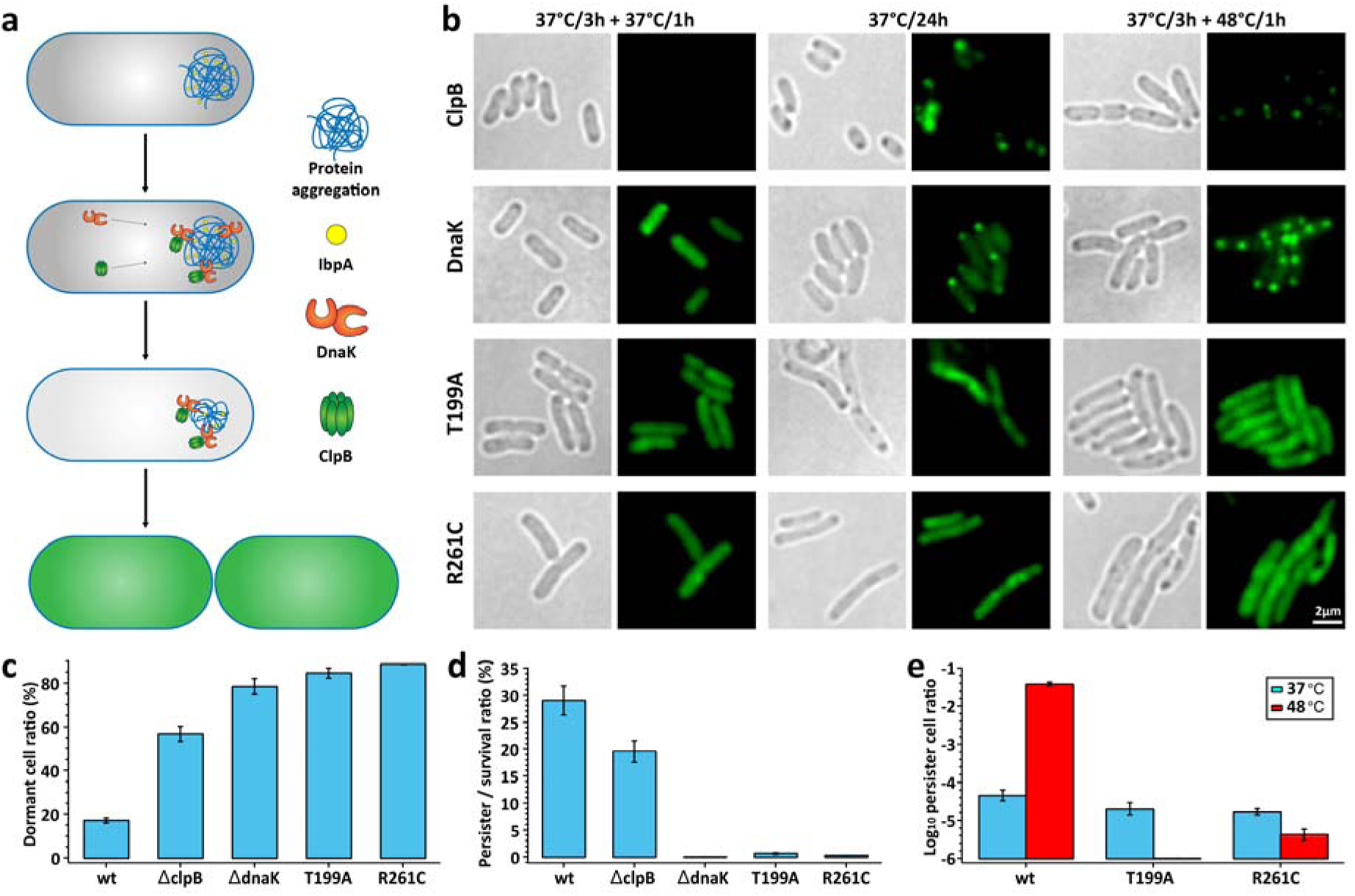
Protein disaggregation is an ATP-dependent process. a, Model of protein disaggregation in persister cells before resuscitation. b, Brightfield and fluorescence images of cells in normal (left), late stationary (middle) and heat shock (right) conditions. ClpB ⸬ EGFP and DnaK ⸬ EGFP foci co-localize with protein aggregates in both late stationary and heat shock conditions (upper two panels); ATPase mutants of DnaK, T199A and R261C, lost the ability to form functional DnaK-ClpB clusters (lower two panels), (Scale bar, 2 μm). c, Dormant cell ratio of cells from different mutants. d, Persister cell / survival cell ratio of different mutants. e, Frequency of persister formation for different mutants strains without (blue) or with (red) heat shock treatment. All errors are standard errors on the mean.

Previous studies have shown that protein disaggregation by DnaK is an ATP-dependent process^20,21^. We constructed two ATPase activity dysfunctional point mutants, *T199A* and *R261C*. Neither mutant was able to form foci either after heat shock or in late stationary phase (Fig. 4b), although protein aggregates indicated by dark foci were quite visible. Compared with the wild type strain, the dormant cell rate of *T199A* and *R261C* increased substantially (Fig. 4c), but the resuscitation rate after removal of antibiotic decreased dramatically (Fig. 4d). These results indicate that DnaK with impaired ATPase activity is dysfunctional in protein disaggregation.

We’ve established that heat shock could increase dormant cell rate through intensified protein aggregation (Fig. 2d and 2e), which increases cell survival rates and potentially elevates persister rates after antibiotic treatment. As expected, in the wild type strain, persister rate is substantially increased after heat shock but in DnaK dysfunctional strains, *T199A* and *R261C*, persister rate is considerably decreased. These results suggest that intensified protein aggregates deepen dormancy depth, which helps cells evade antibiotic killing and increase population survival rate but that only cells with functional disaggregation apparatus are capable of efficient resuscitation, the others remaining trapped in a dormant state.

Colony forming unit (CFU) based viability assays, often used in bacterial antibiotic tolerance studies, lead to negligence of the existence of VBNC cells. Here based on time-lapse fluorescent microscopy we establish a bacterial dormancy depth model for a global understanding of bacterial drug tolerance. In this model VBNC cells are in deeper dormancy depth than persister cells, making it more difficult to restore cell physiological conditions and initiate proliferation (Fig. 1b). Our model explains several apparently contradictory results in recent literature, notably studies by Vega et al. and Hu et al. who show indole can both induce and reduce persister cell proliferation^24,25^. Increased persister formation from indole may be due to moderate dormancy depth being induced through the introduction of a stress in the form of indole while decreased persister formation can be attributed to a deep dormancy depth being induced in the cells leading to increased VBNC cell production.

To tackle bacterial antibiotic tolerance, the generally accepted approach is to wake dormant cells in order to increase antibiotic efficiency^26^. In contrast, we propose a method of limiting persister production that encourages a bacterium into a deeper state of dormancy increasing the proportion of VBNC cells incapable of resuscitation or regrowth. In our study, we establish that dynamic protein aggregation regulates cell dormancy depth, which is critical for bacterial drug tolerance. As a result, to keep cells in deeper dormancy states we can either intensify protein aggregation to an irreversible level or interrupt disaggregation pathways that prevent cells escaping from their dormant state.

## Materials and Methods

### Bacterial strains and culture conditions

Wild type *E. coli* strains MG1655 and BW25113 were purchased from Yale Genetic Stock Center. The *ΔclpB* knockout strain was taken from the Keio Collection^27^. Other strains used in this study were constructed in our lab as follows (Extended Data Fig. 2).

Strains containing chromosomal gene X-fluorescent protein (FP), translational fusion or single gene X knockout mutants were constructed by *λ* red-mediated gene replacement. For fluorescent protein fusion strains^28^: the targeted fluorescent protein (EGFP, TagBFP, or mMaple3) fragment was amplified and inserted to replace the stop cassette of the selection-counter-selection template plasmid (pSCS3V31C). Then the linker-FP-Toxin-CmR fragment was amplified from the template plasmid with homology arm complementary to the flanking sequences of the insertion site on the chromosome, before being transformed into electrocompetent cells with induced recombineering helper plasmid (pSIM6). After 3-5 hours recovery, transformed cells were plated on selection plates containing chloramphenicol. The Toxin-CmR cassette was then removed from the chromosome by another round of *λ* red mediated recombination using a counter selection template. Finally, cells were plated on counter selection plates containing Rhamnose to activate toxin^29^.

For TC-tag (CCPGCC) strains and point mutation strains we amplified the Toxin-CmR cassette from pSCS3V31C to insert into the TC-tag or point mutation target site on the chromosome. The counter selection template containing TC-tag/mutation sequence and flanking chromosome sequence as the homologous arm was used to replace the Toxin-CmR cassette with the TC-tag/mutation sequence. Lon-TCtag and hslU-TCtag strains were obtained by inserting the TC-tag into amino acid position 731 of Lon and position 363 of HslU. Dnak point mutants T199A and R261C were wild type DnaK with amino acid T->A at position 199 and R->C at position 261, respectively. For *ΔdnaK* strain, we used the Keio cassette flanked by FRT sites in the Keio collection to replace the *dnaK* gene on the chromosome. All primers we have used are listed in Supplementary Table 2.

Unless otherwise stated, all strains were cultured in LB broth.

### Fluorescence microscopy

Time-lapse fluorescent imaging was performed on an inverted microscope (Zeiss Observer.Z1)^30^ Illumination was provided by different solid state lasers (Coherent), 405nm for TagBFP, 488nm for FlAsH and EGFP, and 561nm for propidium iodide (PI), respectively. The fluorescent signal of cells was collected with an EMCCD camera (Photometrics Evolve 512). Appropriate filter sets were selected for each fluorophore according to its spectrum.

For FlAsH staining^31^, cells were harvested and washed three times with PBS and treated with 10mM EDTA for 15min to improve membrane permeability. Cells were then re-suspended in PBS buffer supplemented with 8μM FlAsH-EDT2 (Invitrogen) and incubated for 50 min in dark at 37°C.

For Propidium iodide (PI) staining, cells were harvested and washed three times with 0.85% NaCl, and PI dye (Invitrogen Molecular Probes’ LIVE/DEAD *Bac*Light Bacterial Viability Kits) was added to the cells in 0.85% NaCl to a final concentration of 40μM. The cells were incubated for 15min protected from light at room temperature in order to distinguish dead cells from survival cells. The camera exposure time and gain was adjusted to prevent over-saturated image acquisition.

### Time-lapse recording of antibiotic killing and bacteria resuscitation under a microscope

For prolonged bacterial culturing and time-lapse imaging, we used the Flow Cell System FCS2 (Bioptechs)^30^ Cells cultured overnight were collected, washed three times with M9 minimal medium and imaged on a gel-pad containing 2% low melting temperature agarose (volume of cell culture : volume of gel-pad = 1: 20)^32^. The gel-pad was prepared in the center of the FCS2 chamber as a gel island and surrounded by flowing medium buffer with different components according to experimental requirements. The cells were then observed under brightfield/epifluorescence illumination at 37°C.

To record the antibiotic killing and bacteria resuscitation under a microscope, bacterial cells were exposed to 150 μg/ml Ampicillin in 90%M9 + 10%LB medium for 5 hours at 37°C and then fresh growth medium 90%M9 + 10%LB was flushed in. The growth medium was refreshed every 12 hours, allowing cells to recover sufficiently.

### Image analysis and statistics

For the time-lapse fluorescent imaging experiment, the fate of different bacterial cells after treatment with antibiotic was identified. Subsequently, by tracing the pattern and intensity of their fluorescent signal during and after antibiotic treatment we determined the relationship between these characteristics and the cell fate.

Image analysis was performed using ImageJ software (Fiji). Outlines of cells were identified from brightfield images. Pixel intensity of all pixels within a cell were measured from the fluorescent channel after subtraction of the background and auto-fluorescence. From the resulting histogram, peaks in pixel intensity, f, that were above the mean fluorescent intensity of the cell, a, were identified. We defined candidate foci pixels as those which had fluorescent intensity, f>2a. Foci were identified as regions in the cell where more than 4 pixels above this threshold were adjacent to each other.

### Dormant cell ratio and persister/survival ratio determination

LB cultured cells taken from the indicated time points were collected, washed three times with M9 minimal medium and placed on a fresh gel-pad in FCS2 system as described above. To record bacteria resuscitation under a microscope, we followed the procedure described above.

For dormant cell ratio analysis, the samples were spotted on a gel-pad after PI dye staining to exclude dead cells, and their growth was recorded by time-lapse imaging. Dormant cells were defined as cells non-proliferating for at least 6 hours on a fresh gel-pad when observed under a microscope. For persister/survival cell ratio analysis, after 5 hours ampicillin killing in a 37°C shaker, cells were washed to remove antibiotics and stained with PI dye. The cells were observed on a gel-pad to record their recovery under the microscope. Survival cells were distinguished by live/dead bacterial cell staining; persister cells were the survival cells that resumed growth in fresh medium.

### Antibiotic treatment and persister counting assay

Bacterial cultures were diluted by 1:20 into fresh LB containing 150 μg/mL ampicillin and incubated in a shaker (200rpm) for 5 hours at 37°C. Samples were then removed and appropriately diluted in PBS buffer, and spotted on LB agar plates for overnight culturing at 37°C. Colony formation unit (CFU) counting was performed the next day. Extended CFU counting was performed daily for 5 days. CFU experiments were performed in triplicate.

### Heatshock experiment

For the persister counting assay after heat shock treatment, the 16 hour overnight culture was transferred to 1.5 mL tubes and incubated on a dry bath at different temperatures (42°C to 52°C) for 60 min. After heat shock treatment, samples were grown on an LB agarose gel-pad under the microscope for the dormant ratio assay and were diluted by 1:20 into fresh LB with 150 μg/mL ampicillin for persister counting assay.

### Super-resolution fluorescence microscopy and data analysis

Sample preparation, image processing and data analysis was modified from Bates^33,34^. Strain DnaK-mMaple3 was cultured overnight in 4 ml LB at 37°C. Cells were harvested by centrifugation at 5000g for 2 min and washed with filtered PBS 3 times. Cells were then fixed with 4% PFA for 15 min and washed with PBS another 3 times to remove residual PFA. The fixed cells were inserted into a poly-l-lysine coated flow chamber, incubated for 20 min at room temperature and washed with approximately 150 μl (10 times chamber volume) of PBS to remove extra cells. For drift correction and focal plane stability, 100 nm Tetraspeck beads (Invitrogen, T-7279, 1:1000 in PBS) were added to the sample before the chamber was sealed with nail polish.

Cells were imaged using TIRF illumination mode on a custom-built STORM imaging platform based on an Olympus 83 microscope. Three sets of data for each field of view were collected: brightfield, conventional fluorescence (488nm Excitation,5Hz) and super-resolution fluorescence (561nm Excitation). For both brightfield and conventional fluorescence, 10 frames were collected and the images averaged. Super-resolution images were acquired at a frame rate of 60 Hz and the 405 nm laser power was increased to compensate for photo-bleaching of the mMaple3 molecules. Imaging was performed until all the molecules were completely bleached. STORM data was analyzed following the protocol previously described by Bates in 2007^33^.

### Insoluble protein isolation and SDS-PAGE

The method for insoluble protein isolation from bacterial cells was modified from Tomoyasu^35^. Aliquots of bacterial cultures with the same optical density (2 ml of culture of OD_600_=1) were rapidly cooled on ice to 0°C, Cultures were then centrifuged at 5000 g (4°C) for 10 min to harvest the cells. Pellets were suspended in 40 μl buffer A (10mM potassium phosphate buffer, pH 6.5, 1 mM EDTA, 20% (w/v) sucrose, 1 mg/ml lysozyme) and incubated for 30 min on ice. Cell lysate was combined with 360 μl of buffer B (10mM potassium phosphate buffer, pH 6.5, 1 mM EDTA) and sonication while cooling. The pellet fractions were re-suspended in 400 μl of buffer C (buffer B with 2% NP40) to dissolve membrane proteins, and the aggregated proteins were isolated by centrifugation 15 000 g, 30 min, 4°C. NP40-insoluble pellets were washed twice with 400 μl of buffer B, re-suspended in 50 μlof buffer B by brief sonication.

Gel electrophoresis of the aggregated protein was carried out according to published protocols^36^ using 12% SDS ± PAGE and staining with Coomassie brilliant blue.

### Mass spectrometry and data analysis

For protein identification, the Coomassie-stained total aggregated proteins of each sample were cut out of the gel and destained with a solution of 100 mM ammonium bicarbonate in 50% acetonitrile. After dithiothreitol reduction and iodoacetamide alkylation, the proteins were digested with porcine trypsin (Sequencing grade modified; Promega, Madison, WI) overnight at 37 °C^37^. The resulting tryptic peptides were extracted from the gel pieces with 80% acetonitrile, 0.1% formic acid (FA). The samples were dried in a vacuum centrifuge concentrator at 30 °C and resuspended in 10 μl 0.1%FA.

Using an Easy-nLC 1200 system, 5 μl of sample was loaded at a speed of 0.3 μl/min in 0.1% FA onto a trap column (C_18_, Acclaim PepMap ™ 100 75μm x 2cm nanoViper Thermo) and eluted across a fritless analytical resolving column (C_18_, Acclaim PepMap ™ 75 μm x15cm nanoViper RSLC Thermo) with a 75 min gradient of 4% to 30% LC-MS buffer B (LC-MS buffer A includes 0.1% formic acid; LC-MS buffer B includes 0.1% formic acid and 80% ACN) at 300 nl/min.

Peptides were directly injected into a Thermo Orbitrap Fusion Lumos using a nano-electrospray ion source with electrospray voltages of 2.1 kV. Full scan MS spectra were acquired in the Orbitrap mass analyzer (m/z range: 300-1500 Da) with the resolution set to 120,000 (FWHM) at m/z 200 Da. Full scan target was 1×10^6^ with a maximum fill time of 50 ms. All data were acquired in profile mode using positive polarity. MS/MS spectra data were acquired in the Orbitrap with a resolution of 30,000 (FWHM) at m/z 200 Da and higher-collisional dissociation (HCD) MS/MS fragmentation.

The MS data were aligned with *Escherichia coli* Reviewed Swiss-Port database by Proteome Discoverer 2.1 software.

## Acknowledgments

We thank Dr. Xiaowei Zhuang (Harvard University), Dr. Guang Yao (University of Arizona) for valuable discussions. We also thank the Core Facilities at School of Life Sciences, Peking University for assistance with mass spectrometry and we are grateful to Dong Liu for her help with MS data analysis. This work was financially supported by the National Natural Science Foundation of China (No. 31722003, No. 31770925, No. 31370847), and the Recruitment Program of Global Youth Experts to F. Bai., and the National Science and Technology Major Project (No. 2017ZX10304402) to Y. Pu.

## Author Contributions

Y.P., Y.L. designed the study; carried out experiments; Y.P., Y.L., M.Q. and A.M. analyzed the data; T.T. performed mutagenesis experiments; Z.Z. performed super-resolution imaging; F.B. designed and supervised the study; Y.P., Y.L., A.M. and F.B. wrote the manuscript.

